# Exome sequencing identifies ARID2 as a novel tumor suppressor in early-onset sporadic rectal cancer

**DOI:** 10.1101/2020.04.15.040741

**Authors:** Pratyusha Bala, Anurag Kumar Singh, Padmavathi Kavadipula, Viswakalyan Kotapalli, Radhakrishnan Sabarinathan, Murali Dharan Bashyam

**Author notes:** Address for correspondence: Murali Dharan Bashyam, Laboratory of Molecular Oncology, Centre for DNA Fingerprinting and Diagnostics, Uppal, Hyderabad 500039, India; and.

## Abstract

Early-onset sporadic rectal cancer (EOSRC) is a unique and predominant colorectal cancer (CRC) subtype in India. In order to understand the tumorigenic process in EOSRC, we performed whole exome sequencing of 47 microsatellite stable EOSRC samples. Signature 1 was the predominant mutational signature in EOSRC, as previously shown in other CRC exome studies. More importantly, we identified *TP53, KRAS, APC, PIK3R1* and *SMAD4* as significantly mutated (q<0.1) and *ARID1A* and *ARID2* as near-significantly mutated (restricted hypothesis testing; q<0.1) candidate drivers. Unlike the other candidates, the tumorigenic potential of *ARID2*, encoding a component of the SWI/SNF chromatin remodeling complex, is largely unexplored in CRC. shRNA mediated *ARID2* knockdown performed in two different CRC cell lines resulted in significant alterations in transcript levels of cancer-related target genes. More importantly, *ARID2* knockdown promoted several tumorigenic features including cell viability, proliferation, ability to override contact inhibition of growth, and migration besides significantly increasing tumor formation ability in nude mice. The observed gain in tumorigenic features were rescued upon ectopic expression of *ARID2*. Analyses of the TCGA CRC dataset revealed poorer survival in patients with *ARID2* alterations. We therefore propose *ARID2* as a novel tumor suppressor in CRC.

## Introduction

Colorectal cancer (CRC) is the third most common and the fourth largest killer among all cancers worldwide^2^. Based on seminal studies performed mainly in the Western population, up to 80% of CRC is believed to be caused by de-regulated Wnt signaling arising primarily due to somatic inactivation of the *Adenomatous Polyposis Coli (APC)* tumor suppressor gene. As per the CRC progression dogma, subsequent genetic events follow including inactivation of tumor suppressors such as *TP53* and *SMAD4* and activation of oncogenes such as *KRAS/BRAF* and *PIK3CA*^15^. A minor proportion (~15%) of CRC is caused by defective mismatch repair causing ‘microsatellite instability’ (MSI), arising primarily from the somatic bi-allelic DNA methylation based silencing of *MLH1*^12^ and exhibiting significant overlap with a third causal pathway termed CpG Island Methylator Phenotype (CIMP)^47^

Application of next generation sequencing has validated the aforementioned (and additional) mutated genes and altered pathways in CRC. However, majority of these discoveries were based on aging related colon cancer, the major CRC subtype in the Western population. Of note, ethnicity based deviations from the CRC progression dogma are well known in CRC^49^. Thus, focusing on non-canonical CRC subtypes may reveal additional altered CRC genes/pathways. The recent Globocan report reveals India to be ranked second (following China) worldwide for incidence and mortality associated with rectal cancer in younger individuals (Globocan 2018; http://gco.iarc.fr)^4^. More importantly, CRC (especially rectal cancer) incidence in the young has doubled worldwide (cancer incidence in five continents; https://ci5.iarc.fr)^19^. Our previous studies revealed <u>E</u>arly <u>O</u>nset <u>S</u>poradic <u>R</u>ectal <u>C</u>ancer (EOSRC) to be the predominant albeit poorly studied CRC subtype in India possibly driven by non-Wnt tumorigenesis genes/pathways^28, 42^. We now report identification of *ARID2* as a novel tumor suppressor for CRC based on a whole exome sequencing analysis of EOSRC samples.

## Results

### Whole Exome sequencing reveals known and novel characteristics in EOSRC

To identify molecular alterations underlying rectal adenocarcinoma, we performed whole exome sequencing analysis of 47 carefully selected well annotated <u>m</u>icro<u>s</u>atellite <u>s</u>table (MSS) rectal tumor and matched normal sample pairs (EOSRC-IN). All samples were from patients aged below 61 years (average 46 years; range 22-60; Table S1). Sequence data analyses and variant calling (see Methods and Figure S1) identified 17,471 substitutions and 1,432 small insertions and deletions (indels) (Table S2A). The number of substitutions predominated over indels across all samples with a mean rate (per megabase (MB)) of 6.4 (range 1-35 per sample) for substitutions and 0.5 (range 0-2.35 per sample) for indels (Figure 1a). Five samples (EOSRC-IN-1095, 2575, 2603, 2643 and 2669) showed a significantly higher mutation rate (>12/MB) and can be considered to exhibit a ‘Hypermutator-like condition’ as described in The Cancer Genome Atlas (TCGA) study on CRC^6^. Given that all samples were MSS (status of the ‘hypermutator-like’ samples was re-confirmed using the same DNA sample that was used for exome sequencing), it is surprising to find a high mutation rate. The mean rate for substitutions and indels in non-hypermutated (excluding the ‘hypermutator-like’ samples) samples was 4.1 and 0.3, respectively. In comparison, analysis of corresponding TCGA rectal cancer data (MSS, non-POLE, age-matched) revealed a mean mutation rate of 2.89 (range 0.52 – 8.47) for substitutions and 0.03 (range 0 – 0.13) for indels. Similarly, TCGA Colon (MSS, non-POLE, age-matched) showed a mean mutation rate of 3.06 (range 0.52 – 6.16) for substitutions and 0.05 (range 0 – 0.18) for indels.

**Figure 1:**
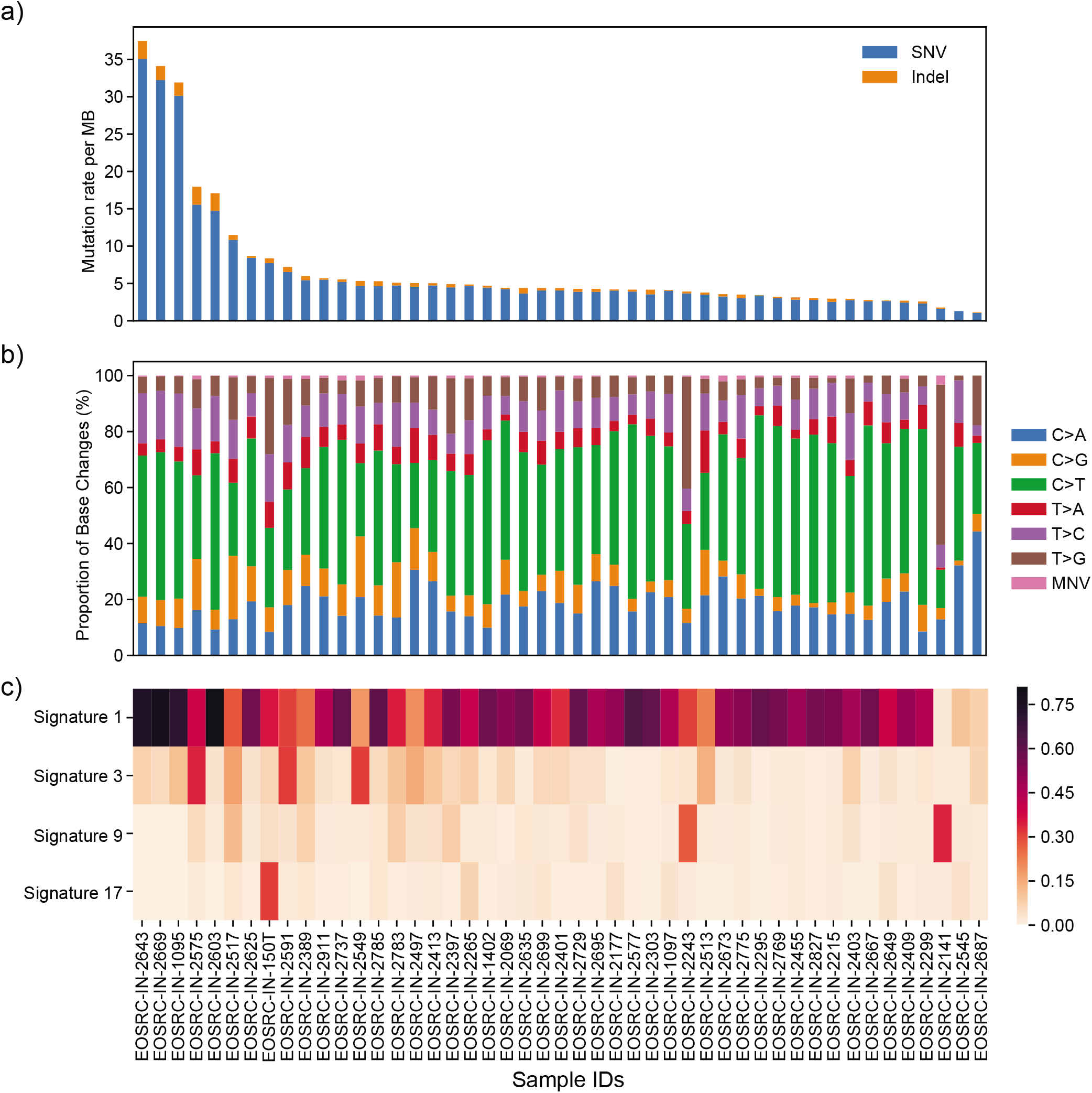
Somatic mutational spectrum in EOSRC. **a)**, Stacked bar plot displaying the mutation rate (mutations per megabase (MB)) separately for SNVs and indels detected in each of the 47 samples. **b)**, Relative contribution of each of the six SNV types as well as of multi nucleotide variants (MNVs) shown as a stacked bar plot. **c)**, Enrichment of COSMIC substitution mutational signatures in each sample. The signatures (1, 17, 3, 9) with higher contribution (i.e., >25% of mutations in a sample can be explained by one of these signatures) is shown here (contribution of all 30 signatures is shown in Figure S2b). The heatmap color indicates the contribution of each signature (colour key is shown on the far right; the scale shows fraction of mutations likely generated by the corresponding mutational signature). Samples are sorted by decreasing mutation rate (from left to right) in each panel.

C>T transitions were the most significant substitution type in most samples (43/47), as reported in previous studies^30^; T>G (EOSRC-IN-2243 and 2141) and C>A (EOSRC-IN-2497 and 2687) transversions abounded in the rest (Figure 1b). More importantly, the overall transition to transversion ratio in our cohort (median 1.3) was significantly lower than the TCGA colon (median 2.4; Mann Whitney U-test *P* = 4.6 × 10^−11^) and rectal (median 2.2; *P* = 6.6 × 10^−7^) tumors (MSS age-matched; see methods) (Figure S2a). Analyses of mutational signatures provides valuable insight into possible genetic and/or environmental causal event(s) driving tumorigenesis. Scrutiny of 30 known COSMIC substitution signatures^1^ in our cohort (using sigfit; see methods) revealed the presence of signatures 1, 3, 9 and 17 (>25% of mutations in a sample could be explained by one of these signatures; Figures 1c and S2b) in EOSRC-IN. Signature 1, associated with the spontaneous deamination of 5-methylcytosine^1^, was detected in almost all samples and is the most frequently reported signature in CRC. Signature 3, associated with defective homologous recombination repair^1^, was identified in three samples (EOSRC-IN-2549, 2575 and 2591) which also showed an increased proportion of longer (> 10 nucleotides; Figure S2c) indels suggesting a possible defective homologous recombination repair in these samples. Signature 9, associated with Polymerase η activity^1^, and Signature 17, likely associated with oxidative damage^1^, were identified in two (EOSRC-IN-2141 and 2243) and one (EOSRC-IN-150T) sample, respectively. No EOSRC-IN sample exhibited signatures (6, 15, 20, 21 or 26)^1^ associated with MSI (figure S2b), as compared to the TCGA MSI group (also analyzed using sigfit; figure S2d), supporting the fact that we selected only MSS samples for the study. In addition, EOSRC-IN samples clustered together with the TCGA MSS samples (which have high Signature 1 exposure), compared to the TCGA hypermutator groups such as POLE mutants and MSI tumors (figure S2d).

### Identification of significantly mutated genes in EOSRC

Of the 18,903 substitutions and indels identified in our cohort, 7,672 variations were predicted to be non-synonymous affecting 5,115 protein-coding genes (Table S2B). The non-synonymous variants included missense (6,803), frameshift (310), stop gained (326), splice site (129), inframe deletion (101), and stop lost (3) (Figures S3a and b). Further, we identified 63 genes (total 590 non-synonymous variants) to be frequently mutated (i.e., non-synonymous mutations found in five or more samples; ~10% of the cohort) (Table S3). We randomly validated 69 of the 590 mutations (~11.5%) using Sanger sequencing with 100% success (Table S3; Figures S3c-d). Of the 63 frequently mutated genes, 13 were previously known to be cancer-associated (COSMIC cancer gene census^45^) (Figure 2a). Of the 13, *TP53* was the most frequently mutated (in 68% of samples) followed by *APC* (45%) and *KRAS* (28% of samples).

**Figure 2:**
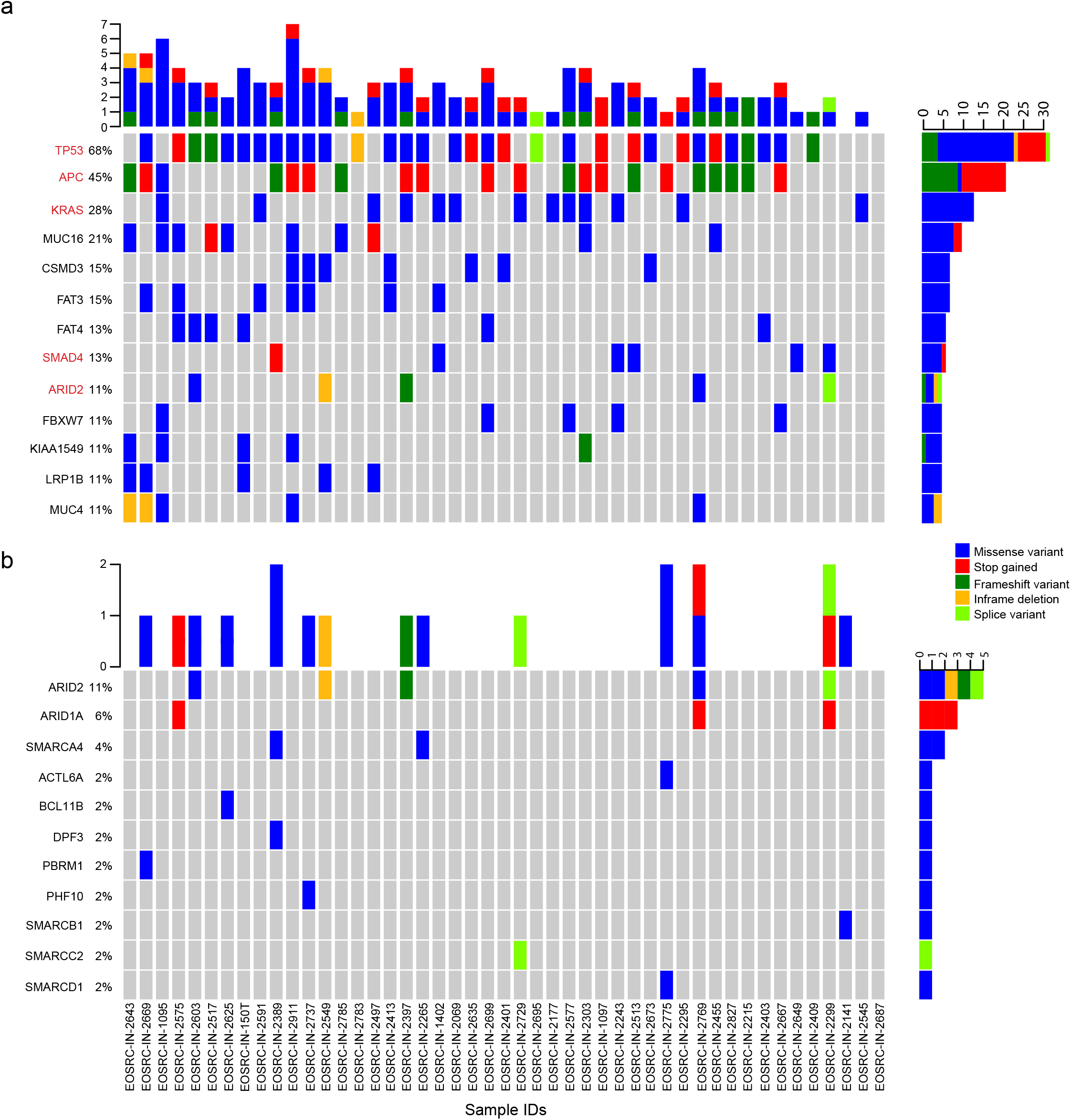
Identification of recurrently mutated cancer-associated genes in EOSRC. **a)**, Frequently mutated (in 5 or more samples) cancer-related genes (COSMIC). The heatmap color indicates the type of non-synonymous mutations observed in a given sample. If a sample has more than one non-synonymous mutation in the same gene, the mutation with high impact (defined based on this order: frameshift, stop gained, inframe deletion, missense and splice variant) was chosen. Genes that were also identified from cancer driver gene detection analysis are indicated in red colour (also see Table S4). **b),** Details of non-synonymous variants identified in various SWI/SNF components in each sample. Mutation types are color coded (indicated), samples are sorted by decreasing mutation rate (from left to right) and percentage of samples harboring mutation in each gene is given on the left; in each panel.

In order to account for gene length and other covariates known to alter mutation frequency, we performed analyses of cancer driver gene detection (samples EOSRC-IN-2643, 2669 and 1095, with mutation rate greater than 30 per MB (hypermutated) were not included in this analysis), based on signals of positive selection analysis (using dNdSCV and oncodriveFML (see methods)), which revealed *TP53, KRAS, APC, PIK3R1, SMAD4* and *ZNF880* as possible driver genes (q<0.1) when all protein-coding genes were considered (Table S4). In addition, *ARID2, ARID1A, TCF12* and *KMT2A* were identified as near-significant candidates (q<0.1) based on the restricted hypothesis testing (using COSMIC cancer gene census^45^, see methods); though only *ARID2* and *ARID1A* were identified using both dNdSCV and oncodriveFML. The frequency of different mutation types for each gene were commensurate with the gene identity; the tumor suppressor *APC* exhibited only inactivating mutations (with one exception) whereas the oncogene *KRAS* exhibited only missense mutations (Figure 2a). *TP53* harbored missense and inactivating mutations in almost equal proportion (Figure 2a) commensurate with its known dual role^26^.

Further, a search for putative ‘hotspot’ mutations^46^ revealed four in *KRAS* of which the frequency of G12V was higher than the other three (G12D, G12S, and G13D) (Table S5A) unlike previous reports which showed G12D to be the most common *KRAS* mutation^40^. We validated the higher frequency of *KRAS* G12V in EOSRC-IN as well as in an extended cohort of 31 EOSRC samples using targeted sequencing (G12V, 14.10%; G12D, 8.97%; G13D, 6.41%; and G12S, 5.13%) (Table S5A). Interestingly, TCGA MSS age-matched rectal cancer samples also exhibited a higher frequency of G12V suggesting it to be a specific feature of EOSRC. In contrast, TCGA colon cancer samples exhibited a higher frequency of G12D (Table S5A).

*TP53* was the most frequently mutated gene in EOSRC commensurate with the high rate of nuclear stabilization detected in our earlier study^42^. *APC* mutations were detected in 21/47 samples (Figure 2a), a frequency of 45%, significantly lower than most previous studies^6, 43^. Majority of somatic tumor mutations in *APC* are restricted to a mutation cluster region (MCR) spanning amino acid positions 1286-1513^38^. We screened the *APC* MCR in 69 EOSRC samples (including the 47 subjected to whole exome sequencing) and detected mutations in 29; a frequency of 42% (Table S5B) thus validating the lower frequency identified in the initial exome screen. The *APC* mutation frequency in three other Asian cohorts (including one from India) were also found to be significantly lower than those reported in studies from the West (Table S5B); whereas there was no discernable difference in *KRAS* and *TP53* mutation frequencies (Table S5B). As per the CRC progression dogma, co-occurrence of mutations in *APC*, *KRAS* and *TP53* is a common feature in CRC^15^. Surprisingly, the mutation co-occurrence frequency in the current study as well as in two other studies from Asia (including one from India) were significantly lower than that detected in studies from the West (Table S5C).

The tumorigenic potential and functional relevance (oncogene or tumor suppressor activity) of *TP53, KRAS, APC, PIK3R1, SMAD4* and *ARID1A* have been extensively studied in CRC^48^. *ARID2* is a subunit of the PBAF SWI/SNF chromatin remodeling complex^51^. Since a) SWI/SNF is characterized as a tumor suppressive complex, b) we identified mutation in a SWI/SNF component in 30% (14/47) of samples (Figure 2b) equivalent or higher than previous estimates^23^ and c) *ARID2* was predicted to be a potential cancer driver (by dNdScv and OncodriveFML) in EOSRC (Table S4), we proceeded to evaluate whether *ARID2* could be a tumor suppressor in CRC.

### ARID2 is a tumor suppressor in CRC

We generated shRNA mediated stable knockdown of *ARID2* in CRC cell lines HCT116 and HT29 (Figure 3a) which resulted in significant alteration of known (cancer-related) *ARID2* target genes including *CCND1, CCNE1, CDKN1B*^13^ and *BMP4*^50^ (figure 3b). More importantly, *ARID2* knockdown caused significantly increased cell growth and viability as measured by BrdU incorporation (Figure 3c) and MTT (Figure 3d) assays, respectively. In addition, reduced *ARID2* expression enabled cells to override contact inhibition of growth (Figure 3e) and exhibit an increased migratory potential as determined through transwell migration assays (Figure 3f), hallmarks of transformed cells. The specificity of effect of *ARID2* was confirmed by its reexpression in stable knockdown cells (Figure 3g) which resulted in rescue of the original phenotype (Figures 3h-i). These results provided strong evidence for a possible tumor suppressor role of *ARID2* in CRC.

**Figure 3:**
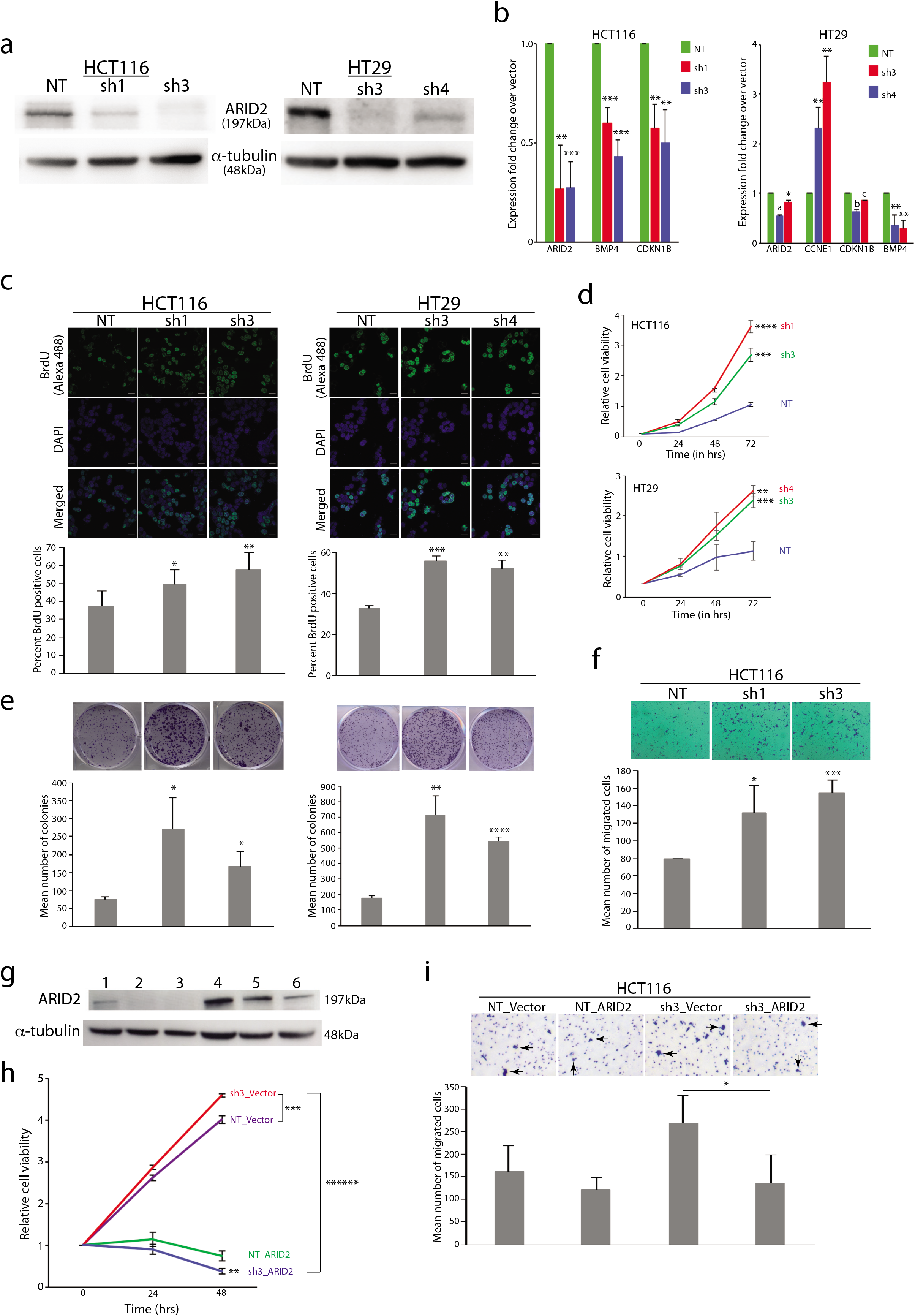
*ARID2* exhibits tumor suppressor activity in HCT116 and HT29 CRC cell lines. **a),** Confirmation of *ARID2* knockdown in HCT116 and HT29 cells using immunoblotting. **b),** *ARID2* knockdown results in alteration of mRNA levels of known transcriptional targets. **c-f),** Various tumorigenic assays reveal possible tumor suppressor activity of ARID2 in CRC cells. **c),** BrdU incorporation assay (scale bars are 20 μm for all fluorescent images); **d)**, cell viability (MTT) assay; **e)**, liquid colony formation assay; **f)**, transwell migration assay (only for HCT116) performed following shRNA mediated *ARID2* knockdown compared to non-targeted control (NT) knockdown. The arrowheads show migrated cells (the smaller multiple blue dots are pores). **g-i)**, Rescue of phenotypes upon *ARID2* expression in ARID2 knockdown HCT116 cells. **g**, Confirmation of ARID2 expression in knockdown cells. Lanes 1, HCT116_ARID2 knockdown + mod-ARID2-GFP; 2, HCT116_ARID2 knockdown + WT-ARID2-GFP; 3, HCT116_ARID2 knockdown + GFP; 4, HCT116_NT knockdown + mod-ARID2-GFP; 5, HCT116_NT knockdown + WT-ARID2-GFP; 6, HCT116_NT knockdown + GFP. **h-i,** Rescue of reduced cell viability (**h**) and of reduced cell migration; black arrow heads indicate migrated cells (**i**) through ectopic expression of *ARID2* in knockdown cells. sh3_vector, HCT116_ARID2 knockdown + GFP; NT_ Vector, HCT116_NT knockdown + GFP; NT_ARID2, HCT116_NT knockdown + mod-ARID2-GFP; sh3_ARID2, HCT116_ARID2 knockdown + mod-ARID2-GFP. Each value represents the mean from at least three independent experiments. Differences were considered significant at a p value less than 0.05 (*, *P*<0.05; **, *P*<0.01; ***, *P*<0.001; a, *P=* 0.0001; b, *P*=0.00005; c, *P*=0.0000009.).

We next tested tumor formation ability of CRC cells in immune-compromised nude mice. Xenograft studies revealed a significantly increased ability of CRC cells harboring *ARID2* knockdown to form subcutaneous tumors (Figure 4a). A tumor suppressor is expected to exhibit reduced transcript levels in tumor compared to normal samples. Indeed, *ARID2* exhibited significantly reduced transcript levels in rectal tumor vs normal samples as determined by RT-qPCR (Figure 4b); similar observations were made from an independent rectal cancer microarray data set^16^ (Figure 4b). Immunohistochemistry (IHC) based evaluation of ARID2 protein expression in a CRC tissue microarray (TMA) (see methods) revealed loss of expression in 20% of tumors; the loss was significantly higher in EOSRC compared to other CRC subtypes (Figures 4c and Table S1). Analysis of TCGA data revealed a significantly poor overall survival in CRC patients harboring *ARID2* alterations (Figure 4d). Finally, a Pan-Cancer analysis performed using TCGA data revealed multiple types of gene alteration in *ARID2* resulting in reduction in transcript levels (Figure 4e).

**Figure 4:**
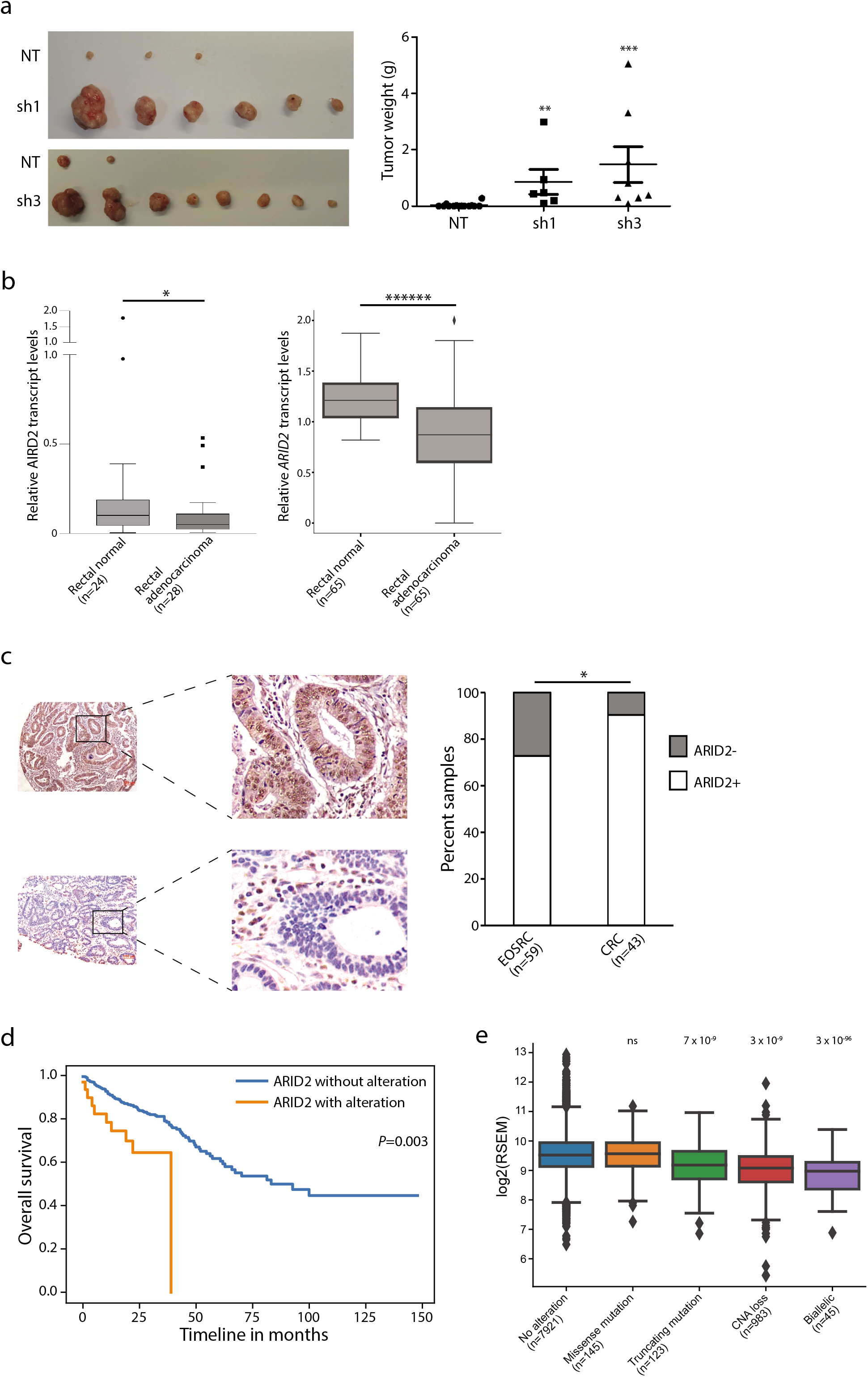
*ARID2* is a tumor suppressor in CRC. **a),** Representative images (left) and quantification of weight of tumor xenografts (right) generated in nude mice with HCT116 cells harboring *ARID2* (sh1 and sh3) or control (NT) knockdown. **b)**, Quantitation of relative *ARID2* transcript levels in tumor vs normal rectal tissues by RT-qPCR in EOSRC-IN (left) as well as from a published microarray gene expression data set (GEO GSE20842) (right). **c**), IHC based determination of ARID2 expression in a CRC TMA. Left panel shows representative images of CRC samples positive (top) or negative (bottom) for ARID2 expression (scale bar of 100μm is shown). Right panel shows bar plot depiction of percentage tumor samples exhibiting loss of ARID2 expression (also see Table S1). ARID2-, ARID2 negative; ARID2+, ARID2 positive. **d)**, Kaplan-Meier analysis of overall survival in CRC patients with and without ARID2 alterations from TCGA data. **e),** Comparison of *ARID2* gene expression levels in TCGA Pan-cancer samples with no *ARID2* alterations or alterations including missense mutation, truncating mutation, copy number loss, or a bi-allelic event. Sample numbers for each alteration type are indicated in brackets. P values, given above each alteration type, denote statistical significance of comparison with no alteration (ns, not significant). Differences were considered significant at a p value less than 0.05 (*, p<0.05; **, p<0.01; ***, p<0.001; ****** p<0.000001).

## Discussion

With several decades of CRC molecular biology research and more than a decade of studies involving targeted and whole exome/genome sequencing, one would expect to have identified all important CRC genes. Nevertheless, by exclusively targeting tumors that a) originated in the rectum, b) were from patients aged 60 years or lower (EOSRC, hugely under-represented in most CRC genomic studies) and c) were from a population shown previously to exhibit deviations from the canonical CRC dogma^28, 29, 42^, we increased the likelihood of identifying a hitherto poorly studied albeit important CRC gene.

Enrichment of mutational signature 1 in EOSRC-IN was commensurate with most other CRC exome/whole genome studies^6^. Surprisingly however, samples EOSRC-IN-2549, 2591 and 2575, exhibited significant enrichment of signature 3 as well as of longer indels, both suggestive of defective homologous recombination repair. The role of Signature 3 in EOSRC needs to be explored further. In addition, five samples (EOSRC-IN-1095, 2575, 2603, 2643 and 2669) exhibited a likely ‘hypermutator’ phenotype (compared to other samples). Given that these samples did not exhibit any known signatures associated with defective DNA repair or replicative polymerases (POLE or POLD), we speculate whether the ‘hypermutator’ like phenotype is driven by a yet unknown somatic mutational process(es).

The p.G12D variant is the most frequent *KRAS* alteration across cancer types including CRC. We surprisingly detected a higher frequency of the p.G12V variant in EOSRC-IN, which was validated in the TCGA data set as well. Of note, previous studies have shown the p.G12V to be associated with a worse prognosis in CRC^21^. The CRC adenoma to carcinoma progression is driven by co-occurring mutations in *APC*, *KRAS* and *TP53;* a well-established model in the Western population^15^. Surprisingly however, a significant minority of samples in our extended cohort as well as in other studies from Asia exhibited co-occurrence of mutations in these three genes. It is interesting to pursue the possibility of an altered sequence of genetic events driving tumor progression in EOSRC in India/Asia. More importantly, *APC* is recommended to be of prognostic value in CRC^43^ but our findings may diminish its utility in EOSRC prognostication and treatment.

The SWI/SNF ATPase subunits BRG1 and BRM, the ARID domain containing subunits ARID1A/B and a few other subunits (PBRM1, INI1, etc.) are frequently mutated in cancers and classified as tumor suppressors^3^. Though potentially inactivating *ARID2* mutations are identified in some cancer types^18, 20, 34^ including MSI+ CRC^5^, a possible tumor suppressor function of *ARID2* is studied only in hepatocellular carcinoma^32^ thus far.

Though we detected possible inactivating mutations in approximately 11% of EOSRC samples, ARID2 loss at protein level was observed in 27% of EOSRC samples (Figure 4c). In addition, TCGA-Pan-Cancer analysis indicated multiple modes of *ARID2* alterations causing a significant reduction in transcript levels (Figure 4e). Taken together, these observations indicate a possible multi-modal *ARID2* inactivation in cancers, which needs to be explored in greater detail. In addition to its well documented role in regulating gene expression^50, 51^, ARID2 plays a dual role in DNA repair by a) interacting with RAD1 to promote homology directed repair^11^ and b) promoting transcriptional repression in double strand break repair^37^. Considering the recent interest to develop targeted therapy against SWI/SNF deficient cancers^44^, it is interesting to determine whether the tumor suppressor role of *ARID2* in CRC is dependent on its transcription regulatory or DNA repair function(s). Given the recalcitrance of advanced colorectal tumors to conventional therapies and the identification of PBAF inactivation association with increased susceptibility to killing by cytotoxic T cells^41^, our identification of *ARID2* loss in a significant proportion of EOSRC, a poorly studied CRC subtype, assumes significance.

## Materials and Methods

### Samples

The study was performed as per the revised Helsinki declaration^7^. Sample collection, storage, processing, isolation of DNA, immunohistochemistry, as well as testing for MSI and Wnt status are described in our previous studies^28, 42^. Clinico-pathological details of samples used for exome sequencing are listed in Table S1. The tumor normal sample DNA pairs were confirmed through standard STR profiling.

### Exome sequencing, data analyses and variant calling

Exome Sequencing of 27 tumor-normal pairs was outsourced to Medgenome Labs Ltd, Bangalore, India. Whole-exome sequencing libraries were prepared using SureSelectXT Human All Exon V5 + UTR kit (75 Mb; Agilent technologies, Santa Clara, CA, USA) as per manufacturer’s instructions and sequenced on HiSeq 2000 to generate paired end 2 × 100bp sequence reads (Illumina, San Diego, USA). Additionally, 20 tumor-normal pairs were sequenced using the Truseq DNA kit (40Mb; (Illumina)) on the NovaSeq (Illumina) to generate paired end 2 × 150bp sequence reads. The data analysis pipeline is shown in Figure S1. Paired-end reads from tumor and matched normal samples were aligned to GRCh38 human reference genome using the Burrows-Wheeler Aligner (BWA) (v0.7.17)^31^ through the Sarek (v2.3) (https://github.com/SciLifeLab/Sarek)^33^ workflow. We achieved an average coverage of 97X and 53X for tumor and normal, respectively, for 27 samples sequenced using the SureSelectXT kit (75 MB; Agilent Technologies, Santa Clara, CA, USA), and 104X and 102X for tumor and normal, respectively, for 20 pairs sequenced using the Trueseq kit (40 MB; Illumina). Further, the GATK toolkit (v4.0.9.0)^36^ was used to mark duplicated reads and to perform base quality score recalibrations. Mutect2^9^ and Strelka2 (v2.9.3) were used independently to call somatic variations (Single Nucleotide Variations (SNVs) and indels). Further, to filter potential germline polymorphisms and sequencing artifacts, we removed somatic variants that overlap with SNPs (with allele frequency > 1%) in gnomAD^24^ and GenomeAsia 100K^17^ or variants in the panel of normals (PoN, which was generated using Mutect2 from all the matched normal samples in our cohort). To reduce low allele frequency variants, we finally selected somatic mutations supported by a minimum of five reads for both reference (in normal) and alternate (in tumour) alleles. 70% of the selected somatic variants were predicted by both callers, and the remaining 23% and 7% were identified by Strelka2 and Mutect2 alone, respectively. These variants were combined and annotated using SnpEff (v4.3^10^. Further, the variants annotated as missense, stop gained, frameshift, inframe deletion, stop lost or splice site, with the predicted functional impact to be high or moderate, were defined as non-synonymous mutations.

### Mutational Signatures and cancer driver gene detection

The exposure of COSMIC mutational signatures (n=30) in each sample was computed using sigfit (v1.3.1) (https://github.com/kgori/sigfit)^22^. OncodriveFML (v2.2.0)^39^ and dNdScv (v0.0.1)^35^ were applied on the somatic variants to detect cancer driver genes with signals of positive selection. The predicted cancer driver genes with q-value less than 0.1 were deemed significant. Further, to enrich for cancer-related genes, we selected genes overlapping with the list of known cancer genes (from COSMIC)^45^ and performed the multiple-testing correction (restricted hypothesis testing) using Benjamini-Hochberg procedure (separately for OncodriveFML and dNdScv). The genes with q < 0.1 from this restricted hypothesis testing were considered as near-significant cancer driver genes. Further, the cancer driver genes were scrutinized for any hotspot mutations (i.e. mutations that occurred at a specific site in >=2 samples) within the coding regions.

### Analyses of TCGA and other published data sets

Information on somatic mutations and gene expression data of TCGA Pan-Cancer was obtained from the MC3 somatic mutational calls^14^ and Firebrowse server (http://firebrowse.org), respectively. Information on clinical annotations such as patient’s age and MSI status were obtained from the GDC portal (https://gdc.cancer.gov). Mutation data was available for 10,295 tumors of which 408 and 151 corresponded to colon and rectal tumors. To ensure an accurate comparison, we selected a subset of TCGA colorectal samples that were microsatellite stable (MSS), devoid of POLE mutations, and were from patients aged below 61 years. This resulted in 79 and 47 colon and rectal tumor samples, respectively, which we designated as TCGA MSS age-matched. The exposure of COSMIC mutational signatures (n=30) in each sample was computed using sigfit (v1.3.1). Relative *ARID2* transcript levels in 65 rectal cancer and normal samples were determined from GEO GSE20842.

### Survival Analysis

Overall Survival Kaplan-Meier Estimate was performed using publicly available TCGA data at cBioPortal using default settings^8^.

### Cell culture and manipulations

HCT116 and HT29 CRC cell lines, authenticated via STR profiling and confirmed to be free of mycoplasma contamination, were maintained in Dulbecco’s Modified Eagle’s Medium (DMEM) supplemented with 10% FBS and antibiotic/antimycotic solution (Thermo Fisher Scientific, Waltham, MA, USA). Stable *ARID2* knockdowns in HCT116 and HT29 were obtained using two shRNAs (sh1 and sh3 for HCT116 and sh3 and sh4 for HT29; shRNA sequences are given in Table S6) separately as described before^27^ Cell line manipulations including MTT, colony formation and transwell migration assays were performed as before^25, 27^ For BrDU assay, cells were seeded in four chamber slides for 24 hours followed by incubation with 10 μg/mL 5- Bromo-2’-deoxyuridine (BrdU) (Sigma-Aldrich, Missouri, USA) for 1 hour (HCT116) or 15 minutes (HT29). Cells were then fixed in 4% paraformaldehyde for 15 minutes, incubated with 2M HCl for 30 minutes and stained with mouse monoclonal anti-BrdU antibody (1:50) (BD Biosciences, San Jose, CA, USA) and Alexa fluor-488 conjugated secondary antibody (1:200) (ThermoFisher scientific). Cells were then mounted using Vectashield mounting medium (Vector laboratories Inc., Burlingame, CA, USA) with DAPI and visualized with LSM 700 confocal laser scanning microscope (Carl Zeiss, Oberkochen, Germany) at 63X magnification. The percentage of BrdU incorporation was determined by counting the number of BrdU-positive nuclei among DAPI-stained nuclei in 10 independent microscope fields. A minimum of 300 nuclei were counted for each experiment.

For rescue experiments, full length ARID2 cDNA from pANT7-ARID2-cGST (Clone 303066, DNASU plasmid repository, Arizona, USA) was subcloned in to pDEST-GFP vector (a kind gift from Dr MS Reddy, CDFD, Hyderabad) to generate WT-ARID2-GFP. The sh3 seed region in *ARID2* was mutated (to prevent/reduce degradation of the *ARID2* transcript under influence of shRNA) through site-directed mutagenesis using primer sequence 5’-GCAACACAGTGTGTCGGATTATCT<u>A</u>CG<u>A</u>CAAAGTTATGGGCTGTCCAT-3 ‘ (the altered nucleotides are underlined) to generate mod-ARID2-GFP which was used for all rescue experiments. Reverse transcription quantitative PCR (RT-qPCR) and immunoblotting were performed as described earlier^27, 28^. For immunoblotting, ARID2 (ThermoFisher scientific) and α-tubulin (Sigma-Aldrich) primary antibodies were used at dilutions of 1:5000 and 1:10000, respectively.

The mean of triplicate experiments was plotted with standard deviation represented as error bars for all assays.

### Mice tumor xenografts

All animal experiments were conducted following approval from CDFD institutional animal ethics committee (Protocol number PCD/CDFD/23). 6-7-week old male FOXN1^−/−^ nude mice (6 and 8 mice for sh1 and sh3, respectively) were subcutaneously injected with 2 × 10^6^ HCT116 cells harbouring *ARID2* or control (non-targeting) knock down in either flank, respectively. Mice were sacrificed 6 weeks post injection by CO_2_ euthanasia and tumors were dissected and weighed.

### CRC TMA and IHC

The CRC TMA and IHC have been described earlier^27^ ARID2 antibody (cat # A302-230A; Bethyl Laboratories, Montgomery, USA) was used at a dilution of 1:1000 dilution. The stained sections were scored independently by two pathologists blinded for the study. Cumulative scores of staining intensities (Negative (0); Weak (1); Moderate (2) and Strong (3)) and percentage positive nuclei (<30% (1); 30-60% (2) and >60% (3)) were calculated for evaluating the stained sections. A score of 0-2 was considered ARID2 negative and of 3-6 as ARID2 positive. Images were taken using Nikon Eclipse 80i (Nikon corporations, Tokyo, Japan) at 20X magnification.

### Statistics

All data obtained from a minimum of three independent experiments were represented as mean +/− standard deviation. The Mann Whitney U test was applied to determine statistical significance of transition/transversion ratio in EOSRC-IN vs TCGA data, nude mice experiments and RT-qPCR experiments; the log rank test for the Kaplan-Meier survival analysis and the Wilcoxon rank sum test for assessing *ARID2* transcript levels. The unpaired student’s t test was applied for all other data.

## Acknowledgements

We thank the patients for kindly agreeing to be a part of the study. We are grateful to Dr Swarnalata Gowrishankar, Apollo hospitals, Hyderabad and Dr Satish Rao, Krishna Institute of Medical Sciences, Hyderabad for evaluation of ARID2 IHC stains on CRC TMA. The work was supported by a grant (SB/SO/HS-007/2013) from the Department of Science and Technology, Government of India to MDB. PB, a registered PhD student of Manipal Academy of Higher Education, is grateful to the University Grants Commission, Government of India for junior and senior research fellowships. We acknowledge CDFD’s Sophisticated Equipment Facility for fluorescence microscopy and Sanger sequencing and the Experimental Animal Facility, for nude mice experiments. RS acknowledges funding support from Ramanujan fellowship.

## Competing interest statement

all authors declare no conflict of interest.

## Author contributions

MDB conceived the project, arranged funding and wrote the manuscript. MDB and RS supervised the research. MDB, PB, RS and AKS designed the experiments and analyzed the results. PB performed experiments involving DNA isolation, QC, Sanger validation, ARID2 stable knockdown, RT-qPCR, and contributed to manuscript writing. PB and PK performed functional assays. Experiments related to construction of the TMA and IHC were performed by VK. AKS and RS performed exome sequencing data analyses and contributed to manuscript writing. All authors contributed to manuscript correction.

## Supplementary Figure Legends

**Figure S1. Data analysis pipeline.** The figure shows steps involved in the processing of exome data and somatic variant calling.

**Figure S2: Analysis of specific sequence variation features and mutational signatures in EOSRC. a),** Distribution of the ratio of transition versus transversion substitutions per sample in three distinct cohorts: EOSRC-IN, TCGA MSS age-matched rectal (READ) and colon (COAD) adenocarcinoma is shown. **b),** Analysis of mutational signatures in EOSRC-IN. The heatmap shows enrichment of 30 COSMIC signatures in each sample of EOSRC-IN; the figure is identical to Figure 1c but shows data for all signatures. **c),** Distribution of the ratio of long (size > 10 bp) versus total indels in each sample of EOSRC-IN. Note the significantly higher ratio for samples (EOSRC-IN-2549, 2575 and 2591) exhibiting enrichment for mutational signature 3 (also see Figure 1c). **d),** The heatmap shows the enrichment of 30 COSMIC signatures in each sample of EOSRC-IN as well as the TCGA CRC cohorts. The color code for each cohort is shown on top. The color of the heatmap indicates the proportion of mutations contributed by each signature in that sample.

**Figure S3: Analyses and validation of mutations identified in EOSRC-IN. a),** Non-synonymous mutation types identified in EOSRC-IN shown as a pie chart. The number and percentage of each non-synonymous mutation type is shown in brackets. **b),** Bar diagram depiction of non-synonymous mutation types identified in each sample. **c-d),** Validation of specific *APC* and *KRAS* (panel c) and *ARID2* (panel d) mutations. In each panel, IGV image (top) as well as electropherogram of the validation through Sanger sequencing (bottom) are shown. The IGV and Sanger sequence for both *KRAS* mutations are for the non-coding strand. The electropherogram result of Sanger sequencing for *ARID2* mutations represent sequence of PCR product cloned into plasmid vectors (and not direct sequence of PCR product); only the mutant sequence is shown.

## Notes

### Competing Interest Statement

The authors have declared no competing interest.

## References

1 Alexandrov LB, Nik-Zainal S, Wedge DC, Aparicio SA, Behjati S, Biankin AV et al. Signatures of mutational processes in human cancer. Nature 2013; 500: 415–421.

2 Arnold M, Sierra MS, Laversanne M, Soerjomataram I, Jemal A, Bray F. Global patterns and trends in colorectal cancer incidence and mortality. Gut 2017; 66: 683–691.

3 Bracken AP, Brien GL, Verrijzer CP. Dangerous liaisons: interplay between SWI/SNF, NuRD, and Polycomb in chromatin regulation and cancer. Genes Dev 2019; 33: 936–959.

4 Bray F, Ferlay J, Soerjomataram I, Siegel RL, Torre LA, Jemal A. Global cancer statistics 2018: GLOBOCAN estimates of incidence and mortality worldwide for 36 cancers in 185 countries. CA Cancer J Clin 2018; 68: 394–424.

5 Cajuso T, Hanninen UA, Kondelin J, Gylfe AE, Tanskanen T, Katainen R et al. Exome sequencing reveals frequent inactivating mutations in ARID1A, ARID1B, ARID2 and ARID4A in microsatellite unstable colorectal cancer. Int J Cancer 2014; 135: 611–623.

6 Cancer Genome Atlas N. Comprehensive molecular characterization of human colon and rectal cancer. Nature 2012; 487: 330–337.

7 Carlson RV, Boyd KM, Webb DJ. The revision of the Declaration of Helsinki: past, present and future. Br J Clin Pharmacol 2004; 57: 695–713.

8 Cerami E, Gao J, Dogrusoz U, Gross BE, Sumer SO, Aksoy BA et al. The cBio cancer genomics portal: an open platform for exploring multidimensional cancer genomics data. Cancer Discov 2012; 2: 401–404.

9 Cibulskis K, Lawrence MS, Carter SL, Sivachenko A, Jaffe D, Sougnez C et al. Sensitive detection of somatic point mutations in impure and heterogeneous cancer samples. Nat Biotechnol 2013; 31: 213–219.

10 Cingolani P, Platts A, Wang le L, Coon M, Nguyen T, Wang L et al. A program for annotating and predicting the effects of single nucleotide polymorphisms, SnpEff: SNPs in the genome of Drosophila melanogaster strain w1118; iso-2; iso-3. Fly (Austin) 2012; 6: 80–92.

11 de Castro RO, Previato L, Goitea V, Felberg A, Guiraldelli MF, Filiberti A et al. The chromatin-remodeling subunit Baf200 promotes homology-directed DNA repair and regulates distinct chromatin-remodeling complexes. J Biol Chem 2017; 292: 8459–8471.

12 de la Chapelle A, Hampel H. Clinical relevance of microsatellite instability in colorectal cancer. J Clin Oncol 2010; 28: 3380–3387.

13 Duan Y, Tian L, Gao Q, Liang L, Zhang W, Yang Y et al. Chromatin remodeling gene ARID2 targets cyclin D1 and cyclin E1 to suppress hepatoma cell progression. Oncotarget 2016; 7: 45863–45875.

14 Ellrott K BM, Saksena G, Covington KR, Kandoth C, Stewart C, et al. Scalable Open Science Approach for Mutation Calling of Tumor Exomes Using Multiple Genomic Pipelines. Cell Syst 2018; 6.

15 Fearon ER, Vogelstein B. A genetic model for colorectal tumorigenesis. Cell 1990; 61: 759–767.

16 Gaedcke J, Grade M, Jung K, Schirmer M, Jo P, Obermeyer C et al. KRAS and BRAF mutations in patients with rectal cancer treated with preoperative chemoradiotherapy. Radiother Oncol 2010; 94: 76–81.

17 GenomeAsia KC. The GenomeAsia 100K Project enables genetic discoveries across Asia. Nature 2019; 576: 106–111.

18 Hodis E, Watson IR, Kryukov GV, Arold ST, Imielinski M, Theurillat JP et al. A landscape of driver mutations in melanoma. Cell 2012; 150: 251–263.

19 Hofseth LJ, Hebert JR, Chanda A, Chen H, Love BL, Pena MM et al. Early-onset colorectal cancer: initial clues and current views. Nat Rev Gastroenterol Hepatol 2020.

20 India Project Team of the International Cancer Genome C. Mutational landscape of gingivo-buccal oral squamous cell carcinoma reveals new recurrently-mutated genes and molecular subgroups. Nat Commun 2013; 4: 2873.

21 Jones RP, Sutton PA, Evans JP, Clifford R, McAvoy A, Lewis J et al. Specific mutations in KRAS codon 12 are associated with worse overall survival in patients with advanced and recurrent colorectal cancer. Br J Cancer 2017; 116: 923–929.

22 K Gori AB-O. sigfit: flexible Bayesian inference of mutational signatures 2018.

23 Kadoch C, Hargreaves DC, Hodges C, Elias L, Ho L, Ranish J et al. Proteomic and bioinformatic analysis of mammalian SWI/SNF complexes identifies extensive roles in human malignancy. Nat Genet 2013; 45: 592–601.

24 Karczewski KJ FL, Tiao G, Cummings BB, Alföldi J, Wang Q, et al. Variation across 141,456 human exomes and genomes reveals the spectrum of loss-of-function intolerance across human protein-coding genes. bioRxiv 2019.

25 Khursheed M, Kolla JN, Kotapalli V, Gupta N, Gowrishankar S, Uppin SG et al. ARID1B, a member of the human SWI/SNF chromatin remodeling complex, exhibits tumour-suppressor activities in pancreatic cancer cell lines. Br J Cancer 2013; 108: 2056–2062.

26 Kim MP, Lozano G. Mutant p53 partners in crime. Cell Death Differ 2018; 25: 161–168.

27 Kumar R, Kotapalli V, Naz A, Gowrishankar S, Rao S, Pollack JR et al. XPNPEP3 is a novel transcriptional target of canonical Wnt/beta-catenin signaling. Genes Chromosomes Cancer 2018; 57:304–310.

28 Kumar R, Raman R, Kotapalli V, Gowrishankar S, Pyne S, Pollack JR et al. Ca(2+)/nuclear factor of activated T cells signaling is enriched in early-onset rectal tumors devoid of canonical Wnt activation. J Mol Med (Berl) 2018; 96: 135–146.

29 Laskar RS, Ghosh SK, Talukdar FR. Rectal cancer profiling identifies distinct subtypes in India based on age at onset, genetic, epigenetic and clinicopathological characteristics. Mol Carcinog 2015; 54: 1786–1795.

30 Lawrence MS, Stojanov P, Polak P, Kryukov GV, Cibulskis K, Sivachenko A et al. Mutational heterogeneity in cancer and the search for new cancer-associated genes. Nature 2013; 499: 214–218.

31 Li H, Durbin R. Fast and accurate short read alignment with Burrows-Wheeler transform. Bioinformatics 2009; 25: 1754–1760.

32 Li M, Zhao H, Zhang X, Wood LD, Anders RA, Choti MA et al. Inactivating mutations of the chromatin remodeling gene ARID2 in hepatocellular carcinoma. Nat Genet 2011; 43: 828–829.

33 M Garcia SJ, M Larsson, PI Olason, M Martin, J Eisfeldt, S DiLorenzo, J Sandgren, TD de Stahl, V Wirtz, M Nister, B Nystedt, M Kaller. Sarek: A portable workflow for whole-genome sequencing analysis of germline and somatic variants. bioRxiv 2018.

34 Manceau G, Letouze E, Guichard C, Didelot A, Cazes A, Corte H et al. Recurrent inactivating mutations of ARID2 in non-small cell lung carcinoma. Int J Cancer 2013; 132: 2217–2221.

35 Martincorena I, Raine KM, Gerstung M, Dawson KJ, Haase K, Van Loo P et al. Universal Patterns of Selection in Cancer and Somatic Tissues. Cell 2018; 173: 1823.

36 McKenna A, Hanna M, Banks E, Sivachenko A, Cibulskis K, Kernytsky A et al. The Genome Analysis Toolkit: a MapReduce framework for analyzing next-generation DNA sequencing data. Genome Res 2010; 20: 1297–1303.

37 Meisenberg C, Pinder SI, Hopkins SR, Wooller SK, Benstead-Hume G, Pearl FMG et al. Repression of Transcription at DNA Breaks Requires Cohesin throughout Interphase and Prevents Genome Instability. Mol Cell 2019; 73: 212–223 e217.

38 Miyoshi Y, Nagase H, Ando H, Horii A, Ichii S, Nakatsuru S et al. Somatic mutations of the APC gene in colorectal tumors: mutation cluster region in the APC gene. Hum Mol Genet 1992; 1: 229–233.

39 Mularoni L, Sabarinathan R, Deu-Pons J, Gonzalez-Perez A, Lopez-Bigas N. OncodriveFML: a general framework to identify coding and non-coding regions with cancer driver mutations. Genome Biol 2016; 17: 128.

40 Neumann J, Zeindl-Eberhart E, Kirchner T, Jung A. Frequency and type of KRAS mutations in routine diagnostic analysis of metastatic colorectal cancer. Pathol Res Pract 2009; 205: 858–862.

41 Pan D, Kobayashi A, Jiang P, Ferrari de Andrade L, Tay RE, Luoma AM et al. A major chromatin regulator determines resistance of tumor cells to T cell-mediated killing. Science 2018; 359: 770–775.

42 Raman R, Kotapalli V, Adduri R, Gowrishankar S, Bashyam L, Chaudhary A et al. Evidence for possible non-canonical pathway(s) driven early-onset colorectal cancer in India. Mol Carcinog 2014; 53 Suppl 1: E181–186.

43 Schell MJ, Yang M, Teer JK, Lo FY, Madan A, Coppola D et al. A multigene mutation classification of 468 colorectal cancers reveals a prognostic role for APC. Nat Commun 2016; 7: 11743.

44 Shorstova T, Marques M, Su J, Johnston J, Kleinman CL, Hamel N et al. SWI/SNF-Compromised Cancers Are Susceptible to Bromodomain Inhibitors. Cancer Res 2019; 79: 2761–2774.

45 Sondka Z, Bamford S, Cole CG, Ward SA, Dunham I, Forbes SA. The COSMIC Cancer Gene Census: describing genetic dysfunction across all human cancers. Nat Rev Cancer 2018; 18: 696–705.

46 Tamborero D, Rubio-Perez C, Deu-Pons J, Schroeder MP, Vivancos A, Rovira A et al. Cancer Genome Interpreter annotates the biological and clinical relevance of tumor alterations. Genome Med 2018; 10: 25.

47 Toyota M, Ahuja N, Ohe-Toyota M, Herman JG, Baylin SB, Issa JP. CpG island methylator phenotype in colorectal cancer. Proc Natl Acad Sci U S A 1999; 96: 8681–8686.

48 Walther A, Johnstone E, Swanton C, Midgley R, Tomlinson I, Kerr D. Genetic prognostic and predictive markers in colorectal cancer. Nat Rev Cancer 2009; 9: 489–499.

49 Wang JY, Hsieh JS, Lu CY, Yu FJ, Wu JY, Chen FM et al. The differentially mutational spectra of the APC, K-ras, and p53 genes in sporadic colorectal cancers from Taiwanese patients. Hepatogastroenterology 2007; 54: 2259–2265.

50 Xu F, Flowers S, Moran E. Essential role of ARID2 protein-containing SWI/SNF complex in tissue-specific gene expression. J Biol Chem 2012; 287: 5033–5041.

51 Yan Z, Cui K, Murray DM, Ling C, Xue Y, Gerstein A et al. PBAF chromatin-remodeling complex requires a novel specificity subunit, BAF200, to regulate expression of selective interferon-responsive genes. Genes Dev 2005; 19: 1662–1667.

